# Increasing growth rate slows adaptation when genotypes compete for diffusing resources

**DOI:** 10.1101/616938

**Authors:** Jeremy M. Chacón, William R. Harcombe

## Abstract

The rate at which a species responds to natural selection is a central predictor of the species’ ability to adapt to environmental change. It is well-known that spatially-structured environments slow the rate of adaptation due to increased intra-genotype competition. Here, we show that this effect magnifies over time as a species becomes better adapted and grows faster. Using a reaction-diffusion model, we demonstrate that growth rates are inextricably coupled with effective spatial scales, such that higher growth rates cause more localized competition. This has two effects: selection requires more generations for beneficial mutations to fix, and spatially-caused genetic drift increases. Together, these effects diminish the value of additional growth rate mutations in structured environments.

**Author Summary:** What determines how quickly a beneficial mutation will spread through a population? The intuitive answer is that mutations that confer faster growth rates will spread at a rate that is relative to the size of the growth-rate benefit. Indeed, this is true in a well-mixed environment where all genotypes compete globally. But most organisms don’t live in a simple well-mixed environment. Many organisms, like bacteria, live in a structured environment, such as on the surface of a solid substrate. Does life on a surface change the expectation about the spread of faster-growing mutants? We developed a mathematical model to answer this question, and found that on a surface, the actual growth rates—not just the relative growth rates—were critical to determining how fast a faster-growing mutant spread through a population. When the simulated organisms grew slowly, competition was basically global and a faster-growing mutant could pre-empt resources from far-away competitors. In contrast, when organisms grew more quickly, competition became much more localized, and the faster-growing mutant could only steal resources from neighboring competitors. This result means that there are diminishing returns to series of mutations which confer growth-rate benefits. This idea will help us predict and understand future and past evolutionary trajectories.

## Introduction

Species can adapt to environmental change through the fixation of beneficial alleles. If the rate of fixation is too low, species may face consequences such as extinction in changing environments (1,2). Life in a spatially-structured habit is generally thought to slow the rate of adaptation (3–6). This is because in spatially-structured environments, competitive interactions are more likely to be localized, and so individuals with a beneficial mutation more often compete with themselves than with ancestors (4). However, even in the presence of spatial structure, slow resource acquisition rates can result in population dynamics which resemble a well-mixed system (7). Therefore, it is uncertain whether the effect of spatial structure on the rate of adaptation is constant or dependent on the biological and environmental details of the species examined.

Whether rare genotypes are benefited or harmed by spatial structure is context-dependent. For example, a rare allelopathic genotype can benefit from spatial structure when it is surrounded by susceptible competitors (8,9). However, this benefit of structure can become a detriment if reduction in founder density puts allelopaths too far from susceptible targets or too close to cheaters (9,10). When spatial structure increases the degree to which competition is just with neighbors (i.e. increases the competition localization), it can compound other mechanisms which affect the rate of adaptation. For example, epistatic interactions between mutations can result in reduced rates of adaptation that diminish the effect of additional mutations (11–14), and these diminishing returns can be compounded by spatial structure (2). Similarly, spatial structure can increase genetic drift and founder effects, thereby reducing the rate of adaptation (7,15). However, neither of these negative effects of spatial structure on the rate of adaptation are universal: they can be ameliorated by increased dispersal rates or resource diffusion, which serve to make a system more well-mixed (4,16,17). Therefore, to predict the evolution of species in spatially-structured environments, it is imperative to quantitatively understand what variables results in an environment with high competition localization, and the causal consequences of such localization on the rate of adaptation.

Here we test how growth rate affects competition localization in a spatially-structured environment, and what the consequences of this are on the rate of adaptation. We hypothesized that, due to the localizing effect that high resource uptake has on competitive interactions (7), higher basal growth rates will require more generations for beneficial mutations to fix.

## Results

We used simulations to test how the absolute growth rate of a population influences the time required for invasion by a mutant with a 10% increase in growth rate. Simulations were run in environments that were either well-mixed or spatially-structured on a torus. Each transfer began with 49 founders. The starting transfer (transfer 0) began with one founder being the 10% faster-growing mutant. Each transfer’s simulation ran until 99% of the resources were consumed. Then, we applied a bottleneck to the population, selecting 49 new founder cells proportional to final genotype frequency at the end of the previous transfer to seed a fresh environment. Transfers continued until the faster-growing mutant genotype reached 90% frequency. In spatially-structured environments, founder cells were randomly arranged each transfer, with different sets of randomizations used for each replicate simulation. Resources were initially homogeneously distributed, and spread via diffusion as they were consumed (Fig. 1A).

**Figure 1:**
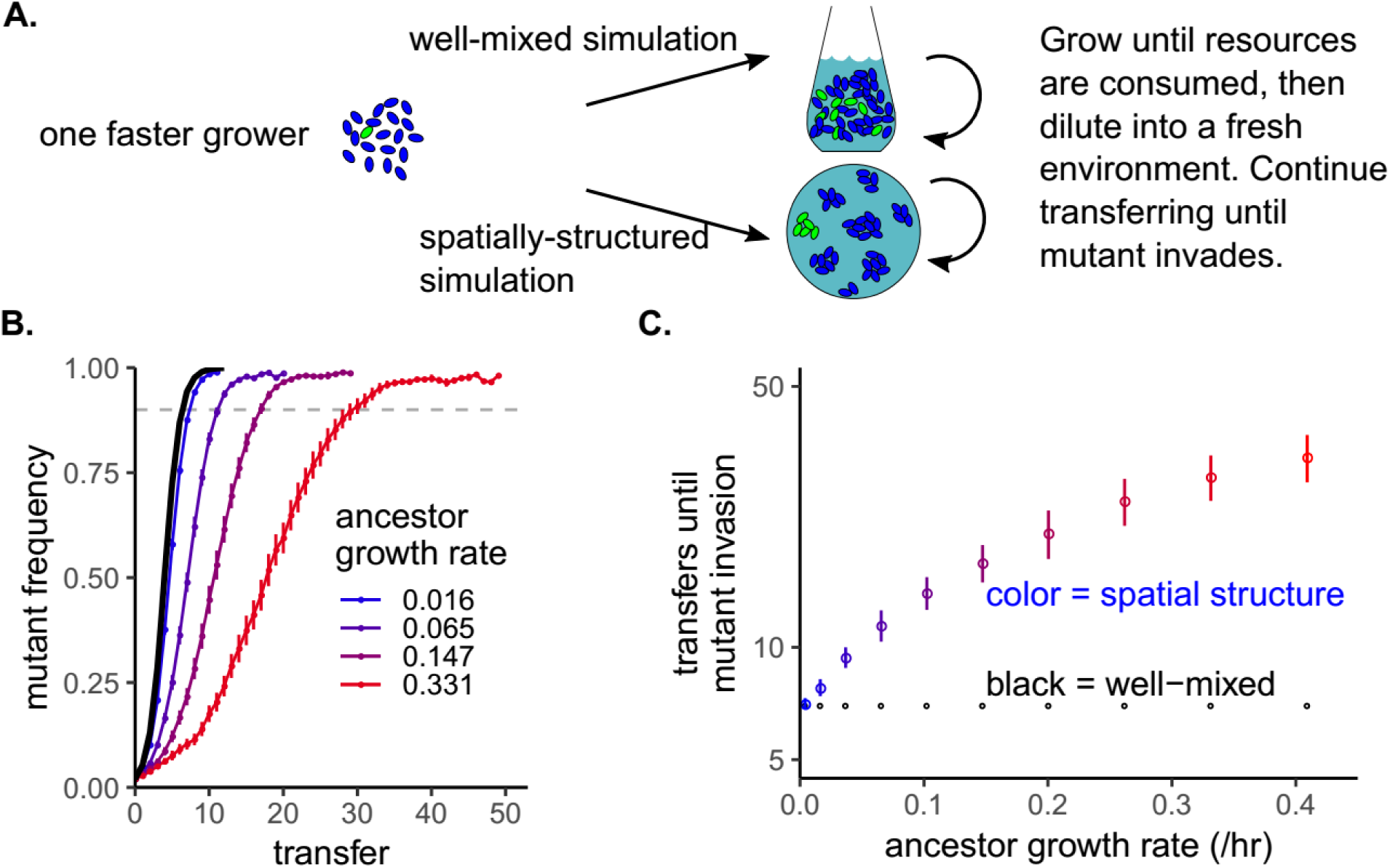
More transfers are required for faster-growing mutants to invade in a spatially-structured environment when their ancestor’s growth rate is higher. A) Schematic for the simulation experiment using resource-explicit models with Monod growth. B) Time series of invasion of faster-growing mutants in simulations, plotted as the mutant frequency versus transfer number. The black line is the time series for all well-mixed simulations (they perfectly overlap regardless of ancestor growth rate). The other colors are for spatially-structured simulations with different ancestor growth rates. Error bars are standard error over twenty replicates. The faster-growing mutant always had a 10% growth rate advantage over its ancestor competitors. The dashed horizontal line indicates the cutoff frequency that was used to determine the number of transfers required for invasion. C) The number of transfers the mutant required to reach a frequency of 0.9, plotted versus ancestor growth rate. Colors correspond to the same simulations in B.

The ancestor’s growth rate had no effect on number of transfers required for the faster-growing mutant to invade a well-mixed environment (black line, Fig. 1B, C). However, in a spatially-structured environment, increasing the ancestor’s growth rate increased the number of transfers required for invasion (Fig. 1B, C). This relationship was robust to changes in the growth rate benefit, the half-saturation constant of resource use, and the number of founder cells. However, the pattern was not observed if resources were replenished as in a chemostat rather than with serial, “seasonal” pulses (Supplementary Fig. 1).

Why did increasing absolute growth rates increase the time required for faster mutants to invade a seasonal spatially-structured habitat? Compared to well-mixed habitats, previous work has shown that the rate of adaptation is slower overall in spatially-structured habitats because interactions are more localized (3–6). We therefore hypothesized that faster ancestor growth rates increased the degree of competition localization. We specifically define “competition localization” as the degree to which one competes with neighbors versus all possible competitors. We sought to test this hypothesis by quantifying competition localization as a function of ancestor growth rate during a single spatial simulation without transfers.

To measure competition localization, we used an approach we previously showed can be used to understand how spatial territory size influences the final biomass of colonies (7). First, the simulation area was converted into a Voronoi diagram (Fig. 2A). This generates a polygon for each founder cell that encloses all of the simulation area that is closer to the focal founder than to any other founder. In other words, the boundaries (and therefore area) of a founder’s polygon are set by its neighbors. For any given founder population, we expect the relationship between polygon areas and final colony sizes to vary when competition is global versus local. If competition is localized, meaning neighbors are most important, then we expect each colony’s final biomass to scale with its polygon area. In contrast, if competition occurs more globally, and adjacent neighbors have proportionally less influence, then polygon area should have a smaller influence on colonies’ final biomasses. By plotting normalized final colony biomasses versus the colonies’ normalized polygon areas and fitting a line to this data, we can measure the degree to which colonies’ biomasses scale with their polygon areas (Fig. 2B). A large slope close to one indicates that neighbors are the most important competitors and therefore competition localization is high. Smaller slopes mean that interactions are more diffuse and competition localization is low. We previously verified this logic by showing that when the slope was close to one, repeating simulations where one founder cell was removed only affected the growth of colonies in neighboring Voronoi polygons (7). Therefore, we deem this slope “competition localization,” and use it as a response variable to test whether competition localization is affected by growth rate.

**Figure 2.**
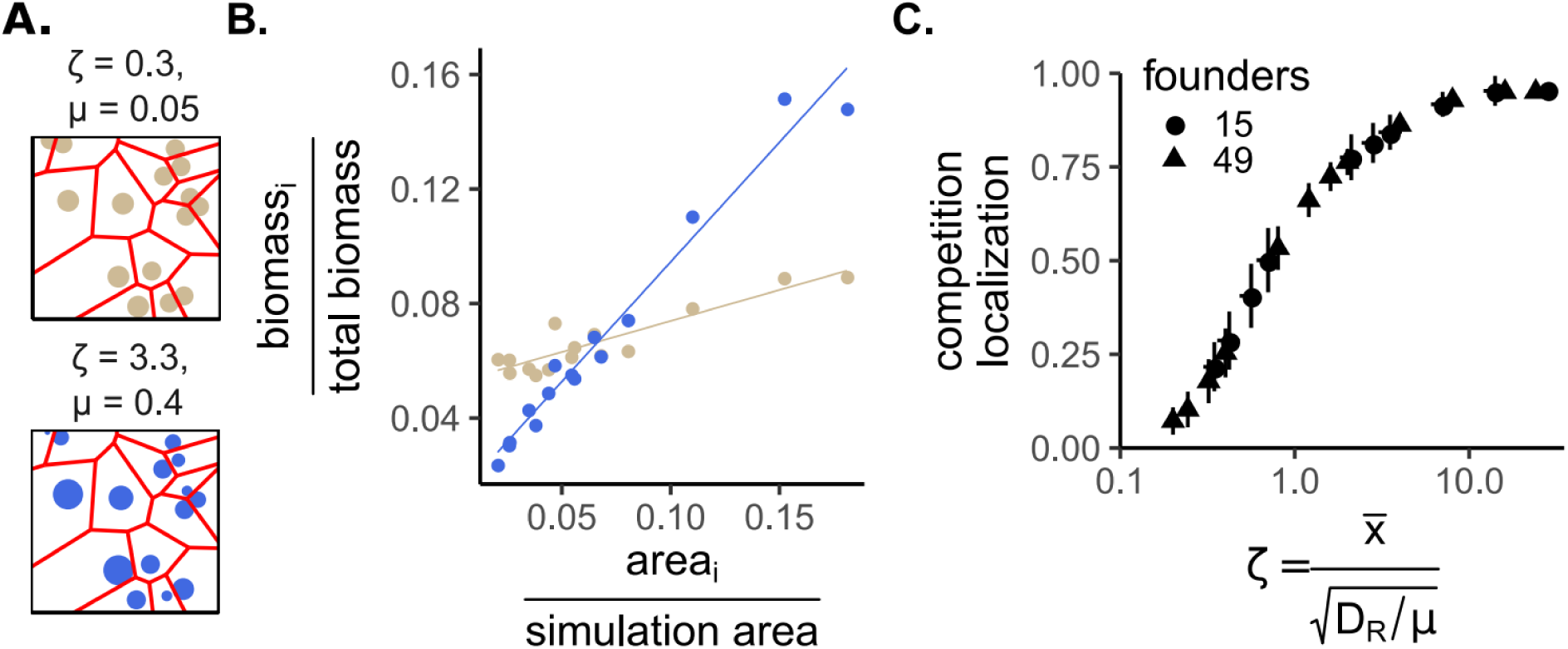
Competition localization increases with growth rate or the distances between colonies, and decreases with the resource diffusion constant. A) Two simulation “maps” with the same founder cell geometry but different parameter values. Circle size is proportional to final biomass of colonies begun with a single founder cell in a spatial environment after resources have been exhausted. The lines designate the Voronoi polygons. The text above each simulation diagram shows 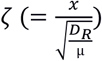 on top, and on the bottom the growth rate (μ) when the resource diffusion constant is similar to that of glucose in agar. B) A plot of relative biomass (biomass of a focal colony divided by the summed biomass of all colonies) versus relative polygon area (polygon area of a focal colony divided by the total simulation area) for the two simulations shown in A. Each point represents a different colony. The line is a linear least-squares fit, and the slope of this line is our measurement of competition localization. C) Competition localization versus ζ for two different founder densities. Vertical error bars are standard error of 20 replicates each with different founder locations. Horizontal error bars are standard error of ζ due to different mean nearest-neighbor distances (see Materials and Methods).

In parallel, we wished to determine whether the rate of mutant invasion was influenced by other parameters coupled to growth rate. This seemed likely as changes in founder density quantitatively (though not qualitatively) altered invasion rates (Supp. Fig. 1). We applied dimensional analysis and scaled the reaction-diffusion model. We found that three variables (including growth rate) could be combined into a single variable. The scaling procedure (see Materials and Methods) showed that

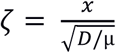

Here, ζ is a scaled parameter combining *D* (the diffusion constant of the resource), µ (the organism’s growth or uptake rate), and *x* (the distance variable). This scaling implies that squared increases in growth rate, or decreases in the resource diffusion constant, are functionally equivalent to linear increases in the distance between competitors. Therefore, simulations can be run under increasing levels of ζ, and interpreted as either increases in inter-competitor distances, increases in growth rate, or decreases in the resource diffusion constant (Supplementary Fig. 2). A similar natural scale was recently observed in a Lotka-Volterra model (18), but its relevance to adaptation or competition localization was not explored.

We used ζ to test the hypothesis that increased competition localization occurs when colonies have a faster growth rate in a spatially-structured environment. Specifically, we ran simulations with increasing levels of ζ (which can be interpreted as increasing levels of growth rate) and measured competition localization using Voronoi diagrams as described above. Most importantly, higher ζ caused increased competition localization (Fig. 2C). Additionally, the method we used to calculate ζ (see Materials and Methods) accurately accounted for differences in founder number / density (Fig. 2C, different shapes).

Increasing ζ, i.e. increasing ancestor growth rate, increased competition localization (Fig. 2) and decreased the rate of adaptation by causing faster-growing mutants to require more transfers to reach high frequency (Fig. 1). We next tested why increasing competition localization decreased the rate of adaptation. Genetic drift can be a result of interactions occurring more locally (15). Therefore, we examined the effect of ζ on the rate of genetic drift among a set of genotypes which had the same growth rate and began at the same genotype frequency. In this scenario, any change in genotype frequency must be due to genetic drift that arose as a result of the stochastic placement of founder cells on the surface. The stochastic nature of initial founder location caused different founders to have different Voronoi polygon areas, which may influence their growth and cause variability in final genotype frequency. We found that the variability in final genotype frequency increased when ζ increased (Fig. 3).

**Figure 3.**
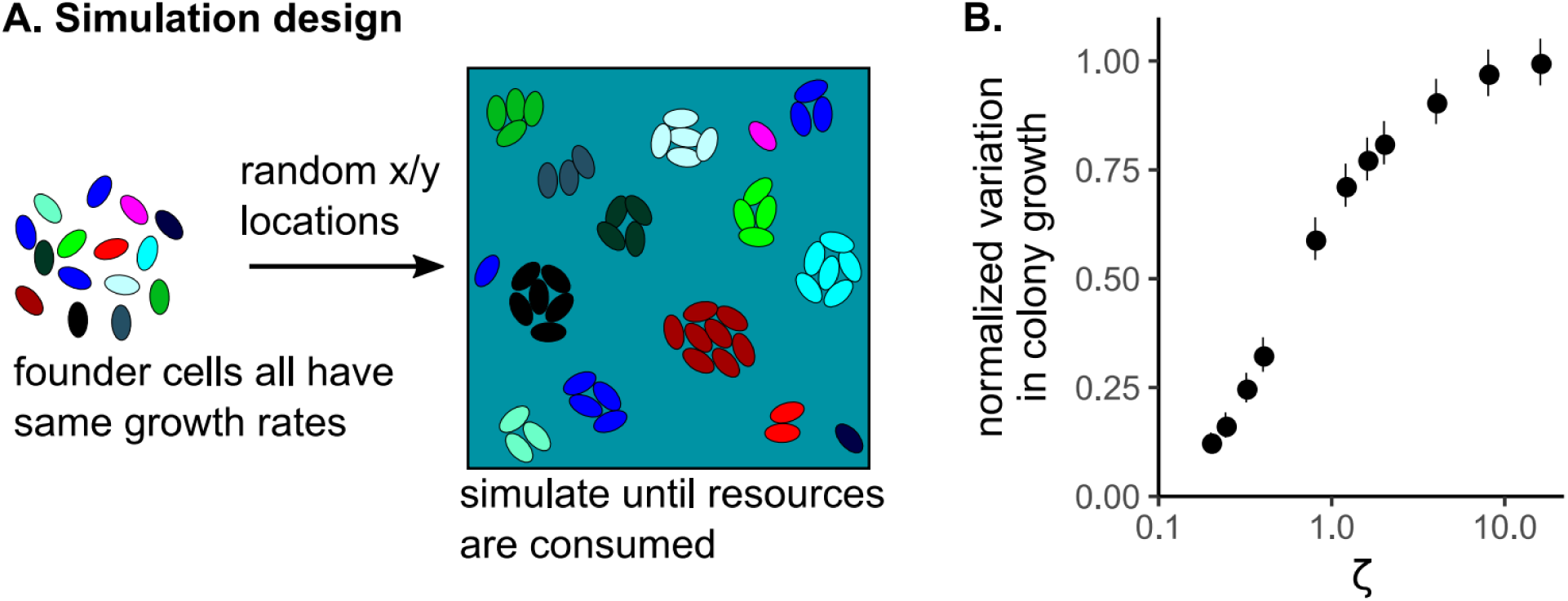
Increasing competition localization increases genetic drift. A) Experimental design for the simulations. B) The variation in frequency of the physiologically-identical genotypes, as a function of ζ, which can be a proxy for growth rate. The points are means, averaged across replicates, of the standard deviation of the final genotype frequency. These means were normalized by dividing by the maximum standard deviation. The error bars are standard error. The horizontal error bars are standard error of the mean of ζ, due to different founder locations causing different mean nearest-neighbor distances (see Materials and Methods).

Finally, we examined how competition localization might affect the rate of selection in an environment without genetic drift. We setup simulations in which founder cells were arranged in a grid with equal inter-colony spacing which removed the stochastic effects of variable Voronoi polygon areas. The ‘center’ cell was a mutant with a 10% growth rate advantage (Fig. 4A). We ran simulations along a gradient of ζ, and measured the change in frequency of the mutant after resources were exhausted. The mutant had a smaller increase in frequency, and therefore was selected for less strongly, when ζ was larger (Fig. 4B).

**Figure 4:**
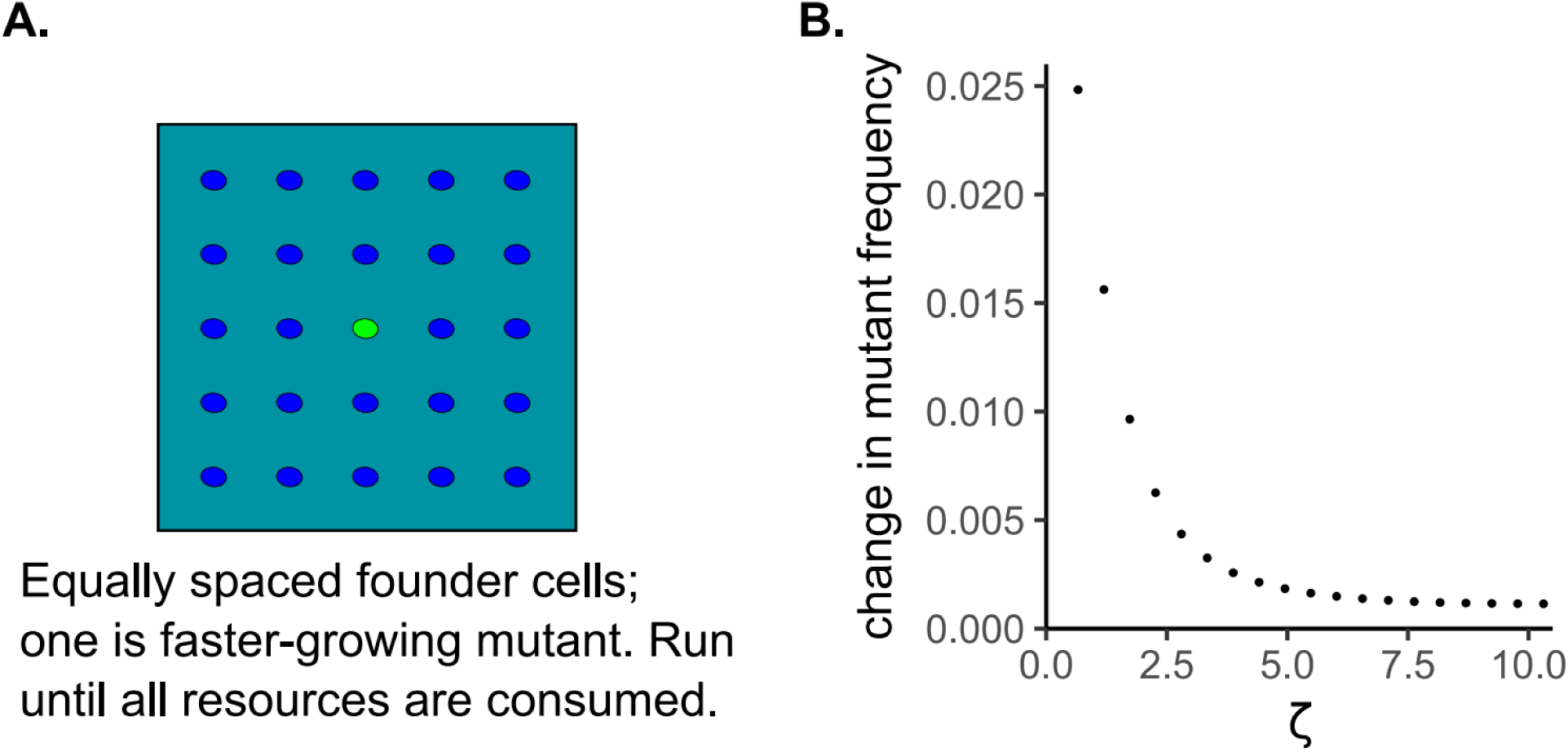
Selection decreases when growth rates are higher in a spatially-structured environment. A) Simulation design with a faster growing mutant (green) in the midst of evenly spaced ancestors. In these simulations, growth rate was varied by varying the scaled parameter ζ. B) The change in frequency of the mutant genotype as a function of ζ.

## Discussion

We showed that higher ancestral growth rates diminish the rate of invasion of a proportionally-faster-growing mutant. The reason for this is that higher basal growth rates cause cells in a spatially-structured environment to compete more locally. The increased competition localization slows selective adaptation generally, because an invading mutant can only preempt resources from neighbors and not more distant competitors. The increased competition localization also increases genetic drift because the size of one’s founding territory becomes a more important determinant of how many offspring a founder will have. Taken together, our results suggest that through evolutionary time increasing growth rate will have progressively diminishing fitness benefits, possibly leading to selection for phenotypes other than growth rate.

We found that in our model, growth rates were inextricably linked to the distances between competing colonies and the diffusion constant of the limiting resource. While we focused on the influence of growth rate, our model shows that we would obtain similar results by reducing the density of founders (thereby increasing inter-competitor distances) or by reducing the diffusion constant of the resource. This result helps explain why reducing nutrient diffusion promotes coexistence of a strong and weak competitor (19). Interestingly, our simulations showed that some “spatially-structured” habitats are functionally equivalent to well-mixed habitats: when growth rates were low enough, spatial position did not alter the outcome of any given founder cell. We expect that continued work which treats growth rates, inter-competitor distances, and resource diffusion rates as interacting parts of one ecosystem property (interaction localization) will help shed light on the sometimes confusing effects of spatial structure.

Our main result on the role of growth rate in reducing the rate of adaptation was robust to many changes in the model: the degree of growth-rate improvement of the invading mutant, the half-saturation resource consumption constant, and the total number of founding cells. However, the result did *not* occur in a chemostat-like environment. Chemostats and batch cultures differ in other ways as well. For instance, chemostats generally select for species which can grow at the lowest resource concentrations (20), whereas batch culture environments select for high maximum growth rates (21,22). Wild environments likely exist in a continuum from seasonal batch culture to pure chemostat, and yet most research focuses on one extreme or the other. We hypothesize that studying model communities in environments that blend aspects of chemostats and batch cultures—as well as better delineating the modes of resource replenishment and mortality in natural systems—will significantly improve our understanding of eco-evolutionary dynamics.

Our simulations used a single limiting resource, with a fixed diffusion constant, in a homogeneous environment. We expect that determining a general version of ζ in more complex situations, for example where species experience co-limitation by multiple resources each with different diffusion constants, will be non-trivial. Nevertheless, we hypothesize that the qualitative result showing that higher growth rates slow adaptation will hold true, as long as the mutant and its ancestors occupy the same niche. However, it is less clear what the outcome will be when colonies alter the diffusion constant of their limiting resource directly, as occurs when resources transition from diffusion in agar to diffusion into the heights of a colony, or when colonies secrete surfactants (23,24). These ecosystem engineering events may change how local the interactions are between colonies, with important implications for the direction and rate of evolution.

Decreasing rates of adaptation to a specific environment are commonly observed (e.g. (25)). The bio-physical cause of diminishing returns shown here is distinct from the diminishing returns due to genetic effects described in well-mixed populations. In well-mixed populations, adaptation can slow because epistatic interactions reduce the benefit of secondary mutations (11,12,26). Here, we showed that diminishing returns result from the fact that increases in growth rate serve to strengthen the effect of spatial structure. High growth rates make it more likely a founder population competes only with neighbors, which decreases the proportion of environment-wide resources the founder consumes, and therefore decreases the founder’s change in allele frequency. In well-mixed populations only relative growth rates matter, while in structured environments absolute growth rates also play a critical role in determining the fate of alleles.

Higher ancestral growth rates reduce selection and are therefore likely to influence evolution of other phenomena. Toxin producers generally gain an advantage from high density because they get more of a return for killing competitors (9). Interestingly, the optimal toxin production rate is predicted to decrease with increases in density because too much production increases the chance that cheaters reap the returns from killed competitors (10). Our results suggest that this relationship will change depending on absolute growth rates, because higher growth rates tend to counteract the effects of higher density. Relatedly, since high growth rates are more likely to restrict interactions to neighbors, we would expect that fast-growing cooperator species are more easily able to exclude cheaters (27).

Considered broadly, our results suggest that absolute growth rate alters the evolutionary trajectory of sessile organisms that compete for diffusing resources. While we modeled bacteria in a homogenous environment, we expect a similar effect in scenarios such as when plants compete for nutrients or water in the soil. Higher growth rates will generally reduce the local pool of resources, thereby reducing selection and increasing genetic drift. The inextricable relationship between growth rate, density and localization of interactions alters fundamental properties of evolutionary trajectories.

## Materials and Methods

All the simulations used a reaction-diffusion partial-differential equations model. Bacteria (B) grew based upon their maximum growth rate (μ), the local resource concentration (R), and Monod kinetics with a half-saturation constant (k). As bacteria grew, resources were consumed with an efficiency parameter (λ). Both bacteria and resources spread via diffusion, with separate diffusion constants (D). We usually simulate two different bacterial genotypes (B_i_), which in the fully-parameterized model each have their own maximum growth rates. The following three equations are the “full” model:

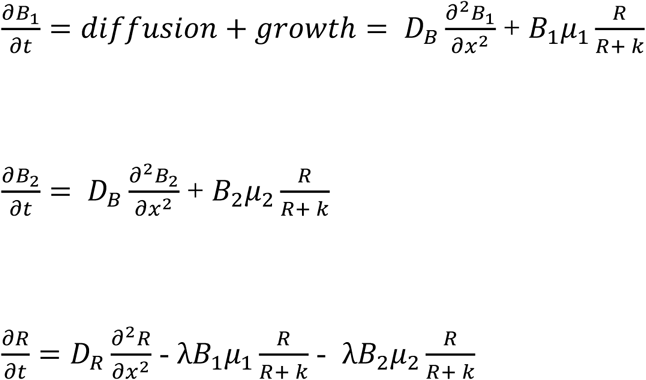

Note that in the above model, x is a vector quantity of [x,y], as all simulations were conducted on planes with equal widths and lengths. We rescaled this model. First, we made state variables dimensionless by division with characteristic quantities of the same dimension:

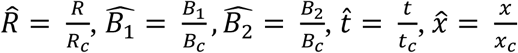

The characteristic variables are defined as:

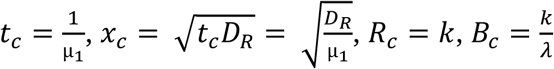

We also redefine the growth rate of the second bacterial genotype based on its proportion relative to the first genotype:

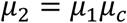

And we redefine the diffusion constant of the bacteria to be proportional to the resource diffusion constant:

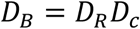

In the model, there is only one distance-related variable, the vector quantity x, which was rescaled to 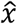.

With the chain rule and some algebra this simplifies the reaction-diffusion model into the following set of equations, which we refer to as the “scaled” model (hats are omitted for clarity):

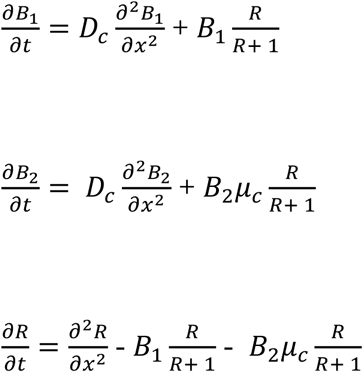

The most interesting result of this scaling, discussed in the Results, is that the variable representing space 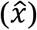 is a function of ‘real’ space (x) as well as bacterial growth rates and resource diffusion:

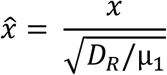

In simulations, this variable is the main representation of space. However, we also use it to predict competition localization by calculating population-level estimates of x from nearest-neighbor distances (see below). To avoid confusion, therefore, when considering how distances, growth rates, and diffusion constants influence different response variables, we rename to 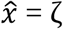 and refer to ζ throughout the manuscript.

The simulations shown in Figure 1 used the full model. For these simulations, a toroidal world with 5 cm per “side” was simulated, discretized into 101 × 101 boxes (i.e. ∼0.05cm / box). D_*R*_ = 5e-6 cm^2^ / s, which is typical for a small sugar in water or a 1% agar Petri dish (7). D_*B*_ = 5e-9 cm^2^ / s, i.e. 1000x smaller than the constant for the resource. k = 1, meaning when there was one cell equivalent of resource left, the growth rate was at half-maximum. Each unit of resource could be converted into one cell, making λ = 1. We also tested k = 50 (Supp Fig. 1A). At the beginning of each transfer, each lattice box contained 100 units of resource. *μ*_2_ (the maximum growth rate of the mutant) was always equal to 1.1*μ*_1_. Bacteria were seeded at 49 randomized locations with one unit of biomass per location. 1 of these founders was the faster-growing mutant. With these boundary conditions, the simulation was run until 99% of all resources were consumed. Once this threshold was reached, the frequency of the faster-growing mutant was calculated. A new environment was generated with new founder locations, and the starting frequency of the bacteria was, allowing for rounding, equal to the final frequency from the previous simulation. These batch-transfer simulations continued until the mutant reached 90% frequency. Twenty replicate batch-transfer experiments were simulated per growth rate, with different founder bacterial locations in each replicate. We also performed equivalent simulations (same founder densities, resource concentrations, growth rates) in mass-action liquid environments.

Simulations testing the relationship between ζ and competition localization (Figure 2) used the scaled model. Each simulation environment used a lattice with 101 × 101 boxes. These simulations used a square (rather than toroidal) lattice, to simplify the spatial analysis. Bacteria were seeded in 49 random locations with 1 cell / box. To track the growth from each founder cell, each founder had its own differential equation. The diffusion constant of the bacteria was 1/1000 that of the resource. As above, at the beginning of the simulation each lattice box contained 100 units of resource. Simulations were run until 99.999% of the resources were consumed, i.e. the bacteria in all simulations went through the same number of generations. Fifteen replicates were simulated per ζ value. Note that ζ is a function of x, the actual physical distance variable. These actual distances, in simulations, are a product of the lattice box size (dx) and the number of boxes between colonies (IC): x = dx * IC. Therefore, we could vary the simulation-level ζ irrespective of the actual founder colony locations by dividing ζ by IC and calculating a 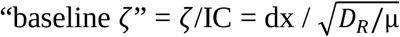. This is the free parameter which we varied in different simulation treatments. However, to get ζ to take into account founder densities, we also had to account for the actual intercolony distances (IC). For any pair of colonies, IC is an exact quantity. But since we were interested in the population-level effect of varying ζ, we estimated the whole-population IC by determining the nearest inter-colony distance for each colony within a simulation, then averaged these to get a population-level average IC in lattice units (Supplementary Figure 3). Multiplying this by “baseline ζ” gives a population-level estimate of ζ. Variations in founder locations, or founder densities, across simulations therefore will cause variations in ζ even for a given lattice box size dx. These variations are shown with the horizontal error bars in Fig. 2C. To test whether this estimate of IC distance was a reasonable approach, in addition to testing many random founder locations for a given “baseline ζ”, we also tested two different founder densities (15 or 49 founder colonies). If our approach was reasonable, the influence of founder density should be subsumed by ζ.

Each replicate had unique founder locations, which were the same for each ζ. Once simulations were complete, competition localization was measured. This was done by measuring the Voronoi polygon area for each founder cell using the dirichletAreas function of the spatstat package in R. Polygon areas were scaled into relative polygon areas by dividing each area by the total simulation area. Relative final biomasses were calculated by dividing the total biomass from each founder cell by the total biomass in the simulation. The competition localization was then calculated by finding the slope of the linear regression of the relative biomass versus the relative polygon areas (Fig. 2B).

Simulations testing the relationship between ζ and the rate of genetic drift (specifically, variance in frequency of neutral genotypes with equivalent growth rates, Fig. 3) used the same simulation results as the simulations testing the relationship between ζ and the Voronoi response.

Simulations examining the influence of ζ on selection when genetic drift cannot have any effect (Fig. 4) used the scaled model with a toroidal lattice of 105 × 105 boxes, each holding 100 units of resource. 49 founder cells were placed at equidistant locations in a grid. One founder had a 10% growth rate advantage. Simulations were run until 99% of the resources were consumed, and the change in frequency of the growth-rate mutant was measured.

All simulations were coded and ran in R. While we initially planned on using the ReacTran package to numerically solve each reaction-diffusion simulation, we found that we could run the simulations much faster and with smaller errors if we iteratively performed each growth step (done per-box using deSolve’s ode function) and diffusion step (using all boxes). Diffusion calculations used a simple forward finite differences scheme, and therefore the time step was kept <= 0.1 * dx^2^ / D, where dx was the width of a box and D was the maximum diffusion constant. This ensured accuracy of the diffusion results. R simulations were run using the University of Minnesota’s Minnesota Supercomputing Institute. Code to run example simulations is provided in Supplementary files 1.

## Acknowledgements

We thank Mike Travisano, members of the Harcombe lab, Nic Vega, and three anonymous reviewers for useful feedback on this manuscript. Funding for this work was provided by the NIH as project #R01GM121498 to William R. Harcombe.

## Competing Interests

The authors have no competing interests.

## Supporting Information

### Compressed file supplementary_code.zip

Contains two files of R code which demonstrate how to run simulations as in the manuscript.

### Supplementary figures

**Supplementary Figure 1.**
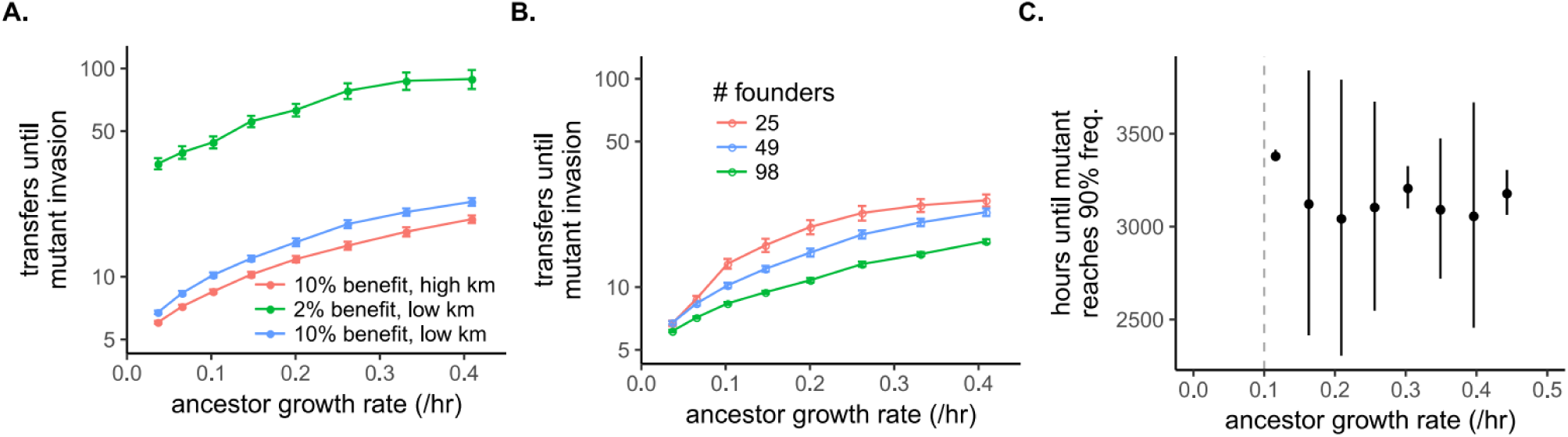
The influence of ancestor growth rate on mutant invasion in a spatially-structured, seasonal environment is robust but does not occur in a chemostat-like environment. A) The number of transfers the mutant required to reach a frequency of 0.9, plotted versus ancestor growth rate. This is similar to Fig. 1C, but includes the results from simulations with either a lower benefit or a higher km. The blue data are the same as from Fig. 1C. B) Like Fig. 1C, but showing the results from simulations with different founder numbers. The simulation size was constant across these treatments, so a higher founder number implies a higher founder density. The blue data are the same as from Fig. 1C. C) The number of hours until a 10% faster-growing mutant genotype reached 90% frequency in a spatial chemostat simulations. The vertical line indicates the chemostat dilution rate (0.1 / hr). Simulations with growth rates below this did not survive and are not plotted. These simulations were similar to those in Fig. 1. A 105×105 box lattice was used to simulate. Boxes each began with 100 resource units. Forty-nine cells were randomly arranged on the lattice. One of these cells was a 10% faster-growing mutant. These simulations different from the main text simulations in that resources were replenished, in each box, from a reservoir with 100 resource units. Additionally, both resources and cells were diluted through time. The dilution rate, which governed replenishment and dilution, was 0.1 / hr. No transfers were performed. Variation in time arose because of the initial random founder placement. Error bars are standard error of 20 replicates.

**Supplementary Figure 2.**
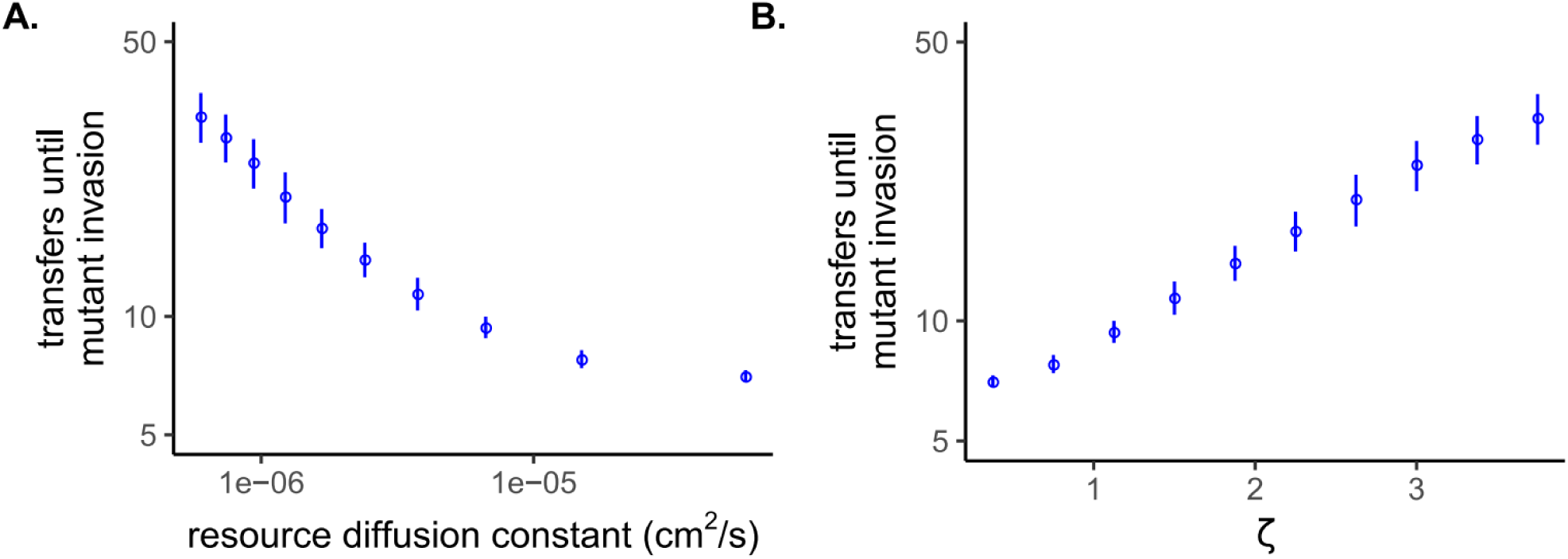
The role of the diffusion constant or zeta on the time required for a faster-growing mutant to invade. The equation for ζ allowed us to consider the results from Fig. 1 as a function of the resource diffusion constant or zeta, rather than as a function of growth rate. A) The number of transfers until the faster-growing mutant reached 90% frequency plotted versus the resource diffusion constant. For this plot, the ancestor growth rate was 0.15 /hr, and the mean inter-colony distance x was 0.03cm. B) The number of transfers until the faster-growing mutant reached 90% frequency plotted versus ζ.

**Supplementary Figure 3.**
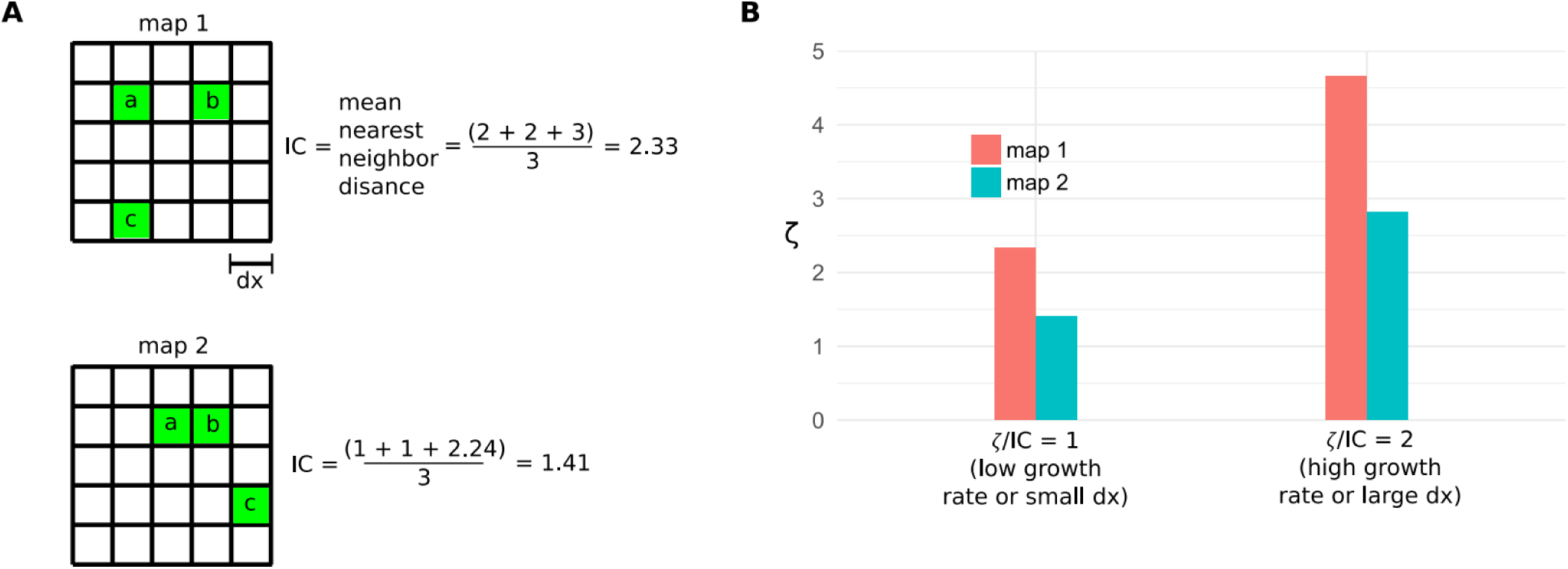
Diagram showing how ζ can change based upon founder locations, even for a given growth rate and lattice box size. A) Two different simplified simulation lattices are shown, with three founder colonies in different boxes. To the right of each map shows how the mean nearest neighbor distance in lattice box units (IC) is calculated. B) ζ plotted versus “baseline ζ” for two different “baseline ζ” and the two different maps in A.

## Notes

#### Summary of Updates

Additional simulations have been added which expand the generality of the main result.

