## supplementary code for "Increasing growth rate slows adaptation when genotypes compete for diffusing resources": simulation_example.r

```
rm(list= ls())
source("./code_for_paper/simulations_functions.r")  # the path here is specific to your system
library(parallel)
library(tidyr)


###########################################
#
# This file provides example code to fun a single
# reaction-diffusion simulation.  The simulations in the main
# text took multiple days to run, using the Minnesota Supercomputer.
# Therefore, here we provide a small example.  If code to reproduce
# the specific simulations from the paper are desired, please contact
# Jeremy Chacon.
#
###########################################


#  This example will simulate growth on a toroidal lattice with two genotypes.
#  The dx(hat) = 0.2 for this simulation, i.e. each lattice box width has a 
#  length value = 0.2.  There will be two genotypes. The mutant genotype has
#  a mu that is 1.1 times that of the the ancestor. There will be 2 mutant founders
#  and 18 ancestor founders.

dxhat = 0.2
genotypes = 2
mu = c(1, 1/1.1) # mutant is the first mu
founders = 20 # total, 2 of these are mutants
mutant_founders = 2

# k and lambda were scaled out, but I use the same function (model_onebox) for 
# the scaled and full models, so I just set them = 1 here
k = rep(1, genotypes) 
lambda = rep(1, genotypes)  

# parms is the list of parameters required by model_onebox
parms = list(species = genotypes,
             mu = mu, k = k, lambda = lambda)

# we will do a small grid for pedagogy; 35 x 35 boxes
N = M = 35 # grid counts in the X and Y directions.

# similar to the parameters k and lambda, Dr has been scaled out, but to
# use model_onebox we set it = 1
Dr = 1 # resource D doesn't exist anymore
Db = rep(Dr/ 1000, genotypes) # bacterial diffusion coef, as fraction of Dr
D = c(Dr, Db)

# each box will start with 100 resource units
resource0 = matrix(100, nrow = N, ncol = M) 
initial_resources = sum(resource0)


# we will randomly locate our founder cells, and make sure they are not overlapping
# initially
bacteria0 = array(0, dim = c(N, M, genotypes)) # diff genotypes get diff z-levels
locs = matrix(sample(1:N, founders * 2, replace = TRUE), ncol = 2)
while(nrow(unique(locs,MARGIN = 1)) != founders){
  locs = matrix(sample(1:N, founders * 2, replace = TRUE), ncol = 2)
  
}
for (i in 1:founders){
  if (i <= mutant_founders){
    bacteria0[locs[i,1], locs[i,2], 1] = 1  # mutant layer
  }else{
    bacteria0[locs[i,1], locs[i,2], 2] = 1  # ancestor layer
  }
}

# take a look:
image(bacteria0[,,1]) # mutant, should be 2 dots
image(bacteria0[,,2]) # ancestor, should be 18 dots

# its much faster to solve the ode system in parallel, so we 
# make a local cluster of 4 processors
cl = makeCluster(4)
# for this to work, we need to tell the cluster to look for the model_onebox function
clusterExport(cl, c("model_onebox"), envir = environment())

# initialize the results, and turn the initial resource and bacteria matrices into one concatenated matrix
results = data.frame()
y0 = abind::abind(resource0,bacteria0, along = 3)


# pick a small enough timestep
dthat = 0.1 * (dxhat * dxhat) / D[1]


# run the simulation loop.  The actual data will be kept in the object "data",
#  which will change between a vector and a matrix depending on whether the ode 
#  or the diffusion is being solved.  The system will run until 99% of all resources
#  are consumed.  We will collect data into the results data.frame every one time unit

# note: this will take about 5 minutes on a modern-ish computer ~ 2018 so be patient!
data = y0
resources_used = 0
results = data.frame()
t_step = 0
while(resources_used < (initial_resources * 0.99)){
  # solve 1 timestep reaction
  data = aperm(parApply(cl,
                        data, c(1,2), single_box, dthat, parms), perm = c(2,3,1))
  
  # this shouldn't happen, but if a NaN occurs, break the simulation
  if (any(is.nan(data))){
    break
    print("NaN found")
  }
  # fix errors (nothing should ever be < 0)--these should be tiny if they occur
  data[data < 0] = 0
  
  # # solve 1 timestep diffusion on a toroidal lattice
  dC = calculate_diffusion_toroid(data, D, dthat, dxhat)
  
  # sum the reaction and diffusion
  data = data + dC
  
  # collect data if one time unit has passed
  t_step = t_step + 1
  if (((t_step * dthat) %% 1) < dthat){
    yields = get_yields(list(data))
    yields$dxhat = dxhat
    yields$time = t_step * dthat
    yields$dthat = dthat
    yields$t_step = t_step
    yields$mu = c(mu, NA)
    resources_used = initial_resources - sum(data[,,1])
    results = rbind(results, yields)
  }
}

# the final biomass maps are still in data
image(data[,,2]) # mutant
image(data[,,3]) # ancestor

# we can view the biomass timeseries results with ggplot
results %>%
  filter(colony != "resource") %>%
  mutate(genotype = ifelse(colony == "2", "mutant","ancestor")) %>%
  ggplot2::ggplot(aes(x = time, y = biomass, color = genotype))+
  geom_line()

# or examine the change in frequency of the mutant
results %>%
  filter(colony != "resource") %>%
  mutate(genotype = ifelse(colony == "2", "mutant","ancestor")) %>%
  group_by(time) %>%
  summarize(mutant_freq = biomass[genotype == "mutant"] / sum(biomass)) %>%
  ggplot2::ggplot(aes(x = time, y = mutant_freq))+
  geom_line()
```
