## supplementary code for "Increasing growth rate slows adaptation when genotypes compete for diffusing resources": simulations_functions.r

```
library(deSolve)
library(abind)
library(dplyr)
library(ggplot2)
library(gridExtra)


#######################################################
# calculate_diffusion_toroid is just like calculate_diffusion,
# but for toroids
#######################################################
calculate_diffusion_toroid = function(data, D, dt, dx){
  n_responses = length(D)
  
  dC = sapply(1:n_responses, function(x) calculate_dC_toroid(data[,,x]), simplify = "array")
  
  # step 2: apply diffusion constants, dt, dx
  dC = sapply(1:n_responses, function(x) dC[,,x] * D[x] * dt / (dx * dx), simplify = "array")
  dC
}

#######################################################
# calculate_diffusion returns the result of applying 
# diffusion to the data with diffusion constants D, timestep dt, 
# and lattice box size dx.  This particular function calculates
# diffusion on rectangle with no-flux edges
#######################################################
calculate_diffusion = function(data, D, dt, dx){
  # this calculates diffusion on a 3d matrix, doing one x-y slice at a time.
  # for example, if you had a resource layer, and a species layer, this
  # would calculate diffusion on each in turn.
  #
  # data is the 3d array, with dim 1 = x, dim 2 = y, dim 3 = the different response variables
  # D is a vector of diffusion constants--must be one per layer in data
  # dt is the time step
  # dx is the grid size
  #
  # the result of this function should be added to the data for the diffusion to occur
  #
  # e.g.
  #  data = reaction(data)
  #  data = data + calculate_diffusion(data, D, dt, dx)
  
  n_responses = length(D)
  
  dC = sapply(1:n_responses, function(x) calculate_dC(data[,,x]), simplify = "array")
  
  # step 2: apply diffusion constants, dt, dx
  dC = sapply(1:n_responses, function(x) dC[,,x] * D[x] * dt / (dx * dx), simplify = "array")
  dC
}

###################################################
# calculate_dC_toroid is just like calculate_dC, but it 
# calculates the forward-finite-differences solution of the laplace
# operator on a toroid
###################################################
calculate_dC_toroid <- function(mat, sanity_check = FALSE){
  dC = matrix(0, nrow = nrow(mat), ncol = ncol(mat))
  
  nrows = nrow(dC)
  ncols = ncol(dC)
  
  # internal diffusion
  dC[2:(nrows-1),2:(ncols-1) ] = mat[1:(nrows-2), 2:(ncols-1)] +
    mat[3:(nrows), 2:(ncols-1)]+
    mat[2:(nrows-1), 1:(ncols-2)]+
    mat[2:(nrows-1), 3:(ncols)]-
    4 * mat[2:(nrows-1),2:(ncols-1) ]
  
  # diffusion on borders (not corners tho)
  # left border
  dC[2:(nrows-1), 1] = mat[1:(nrows-2), 1] +  # shift up
    mat[3:(nrows), 1] +  # shift down
    mat[2:(nrows-1), 2] + # shift right
    mat[2:(nrows-1), ncols] - # shift across-left
    4 * mat[2:(nrows-1), 1] # subtract self
  
  # right border
  dC[2:(nrows-1), ncols] = mat[1:(nrows-2), ncols] +  # shift up
    mat[3:(nrows), ncols] +  # shift down
    mat[2:(nrows-1), ncols - 1] + # shift left
    mat[2:(nrows-1),1] - # shift across right
    4 * mat[2:(nrows-1), ncols] # subtract self
  
  # top border
  dC[1, 2:(ncols-1)] = mat[1, 1:(ncols-2)] +#shift left
    mat[1, 3:(ncols)] + # shift right
    mat[2, 2:(ncols-1)] + # shift down
    mat[nrows,2:(ncols-1)] - # shift across up
    4 * mat[1, 2:(ncols-1)] # subtract self
  
  # bottom border
  dC[nrows, 2:(ncols-1)] = mat[nrows, 1:(ncols-2)] + # shift left
    mat[nrows, 3:(ncols)] + # shift right
    mat[(nrows-1), 2:(ncols-1)] + # shift up
    mat[1, 2:(ncols-1)] - # shift across down
    4 * mat[nrows, 2:(ncols-1)]  
  
  #diffusion on corners
  dC[1,1] = mat[2,1] + mat[1,2] + mat[nrows,1] + mat[1,ncols] - 4 * mat[1,1]
  dC[nrows,1] = mat[nrows-1,1] + mat[nrows,2] + mat[nrows, ncols] + mat[1,1] - 4 * mat[nrows,1]
  dC[1,ncols] = mat[1,ncols-1] + mat[2,ncols] + mat[nrows,ncols] + mat[1, 1] - 4 * mat[1,ncols]
  dC[nrows,ncols] = mat[nrows-1,ncols] + mat[nrows,ncols-1] + mat[1,ncols] + mat[nrows,1] - 4 * mat[nrows,ncols]  
  
  if (sanity_check){
    print(paste("sum(dC) =",sum(dC)))
  }
  dC
}


###################################################
# calculate_dC computes the forwards-finite-differences diffusion calculation
# It assumes thes edges of the simulation lattice are no-flux edges.  
# I've found that it is faster to do this and use a small timestep than it is
# to use the reacTran packages
####################################################
calculate_dC <- function(mat, sanity_check = FALSE){
  dC = matrix(0, nrow = nrow(mat), ncol = ncol(mat))
  
  # internal diffusion
  dC[2:(nrow(dC)-1),2:(ncol(dC)-1) ] = mat[1:(nrow(dC)-2), 2:(ncol(dC)-1)] +
    mat[3:(nrow(dC)), 2:(ncol(dC)-1)]+
    mat[2:(nrow(dC)-1), 1:(ncol(dC)-2)]+
    mat[2:(nrow(dC)-1), 3:(ncol(dC))]-
    4 * mat[2:(nrow(dC)-1),2:(ncol(dC)-1) ]
  
  # diffusion on borders (not corners tho)
  dC[2:(nrow(dC)-1), 1] = mat[1:(nrow(dC)-2), 1] + 
    mat[3:(nrow(dC)), 1] + 
    mat[2:(nrow(dC)-1), 2] -
    3 * mat[2:(nrow(dC)-1), 1]
  
  dC[2:(nrow(dC)-1), ncol(dC)] = mat[1:(nrow(dC)-2), ncol(dC)] + 
    mat[3:(nrow(dC)), ncol(dC)] + 
    mat[2:(nrow(dC)-1), ncol(dC) - 1] -
    3 * mat[2:(nrow(dC)-1), ncol(dC)]
  
  dC[1, 2:(ncol(dC)-1)] = mat[1, 1:(ncol(dC)-2)] +
    mat[1, 3:(ncol(dC))] +
    mat[2, 2:(ncol(dC)-1)] -
    3 * mat[1, 2:(ncol(dC)-1)]
  
  dC[nrow(dC), 2:(ncol(dC)-1)] = mat[nrow(dC), 1:(ncol(dC)-2)] +
    mat[nrow(dC), 3:(ncol(dC))] +
    mat[(nrow(dC)-1), 2:(ncol(dC)-1)] -
    3 * mat[nrow(dC), 2:(ncol(dC)-1)]  
  
  #diffusion on corners
  dC[1,1] = mat[2,1] + mat[1,2] - 2 * mat[1,1]
  dC[nrow(dC),1] = mat[nrow(dC)-1,1] + mat[nrow(dC),2] - 2 * mat[nrow(dC),1]
  dC[1,ncol(dC)] = mat[1,ncol(dC)-1] + mat[2,ncol(dC)] - 2 * mat[1,ncol(dC)]
  dC[nrow(dC),ncol(dC)] = mat[nrow(dC)-1,ncol(dC)] + mat[nrow(dC),ncol(dC)-1] - 2 * mat[nrow(dC),ncol(dC)]  
  
  if (sanity_check){
    print(paste("sum(dC) =",sum(dC)))
  }
  dC
}


########################################################
#  single_box is just a helper function which simplifies 
#  calls to deSolve::ode to solve the system of equations
#  specified by model_onebox
########################################################
single_box = function(y, dt, parms){
  # this is just a helper function, it calls model_onebox
  # on a single vector of data. It's useful for parallelizing the 
  # calls to deSolve::ode
  unname(deSolve::ode(y, c(0, dt), func = model_onebox, parms = parms)[2,2:(length(y)+1)])
}


########################################################
#  model_onebox simulates the dynamics within one lattice box 
#  in between diffusion steps.  It is a function which is used
#  as an argument to a call of deSolve::ode, the numerical solver.
########################################################

model_onebox = function(t,y,parms){
  # parameter grabbing:
  species = parms[[1]] # number of bacterial species 
  mu = parms[[2]] # growth rates
  k = parms[[3]] # half-saturation constants
  lambda = parms[[4]] # resources consumed per bacteria produced
  # state variable grabbing and initialization of derivatives: 
  R = y[1] # resources
  dR = 0 # derivative of R
  B = y[2:length(y)]
  dB = rep(0, species)
  # the actual reaction formulae
  dB = mu * B * R / (R + k)
  dR = sum(-lambda * mu * B * R / (R + k))
  
  list(c(dR,dB)) 
}


###############################################
# get_yields is a helper function which gets a summary
# of the data out of the matrix of the data
##############################################
get_yields = function(data){
  # assumes first layer is resource, all others are colonies
  yields = data.frame()
  for (k in 1:length(data)){
    curr_tp = data[[k]]
    for (j in 2:dim(curr_tp)[3]){
      yield = sum(curr_tp[,,j])
      yields = rbind(yields,
                     data.frame(time = k,
                                colony = j,
                                biomass = yield))
    }
    yields = rbind(yields,
                   data.frame(time = k, colony = "resource",
                              biomass = sum(curr_tp[,,1])))
  }
  yields
}
```
